# M9 Medium Composition Alters *E. coli* Metabolism During Recombinant Expression

**DOI:** 10.1101/2025.07.28.667263

**Authors:** Çağdaş Dağ, Oktay Gocenler, Nilufer Cakir, Merve Turgut, Alp E. Kazar

## Abstract

M9 minimal media and its enhanced variants (M9+ and M9++) are widely used for recombinant protein expression in Escherichia coli, particularly for isotopic labeling required in structural biology techniques such as NMR spectroscopy. This study investigates how different compositions of M9-based media (M9, M9+, and M9++) influence bacterial growth, metabolic stress, and central carbon metabolism during recombinant expression of the protein. Using 1D ^1^H NMR spectroscopy and multivariate statistical analysis, we observed distinct media-dependent metabolic shifts. Standard M9 exhibited limited bacterial growth and heightened stress-related fermentation, indicated by high ethanol and acetate levels. In contrast, M9+ significantly increased biomass but promoted pronounced overflow metabolism. M9++ presented intermediate biomass levels and markedly reduced overflow metabolites, favoring biosynthesis pathways, notably increasing valine, acetoin, and formate concentrations. These findings suggest that further optimization of glucose concentration, nitrogen sources, and phosphate buffering could significantly improve the metabolic balance of M9++, creating an enhanced medium tailored for efficient, high-quality recombinant protein expression and isotopic labeling in *E. coli*.

## Introduction

Recombinant protein production conceptualizes main steps in biochemical and structural biology studies and Escherichia coli is one the most known platforms for the production (Overton, 2014; Terpe, 2006; Nothaft & Szymanski, 2013; Walsh, 2010; Li *et al*., 2025; de Marco, 2025; Wang *et al*., 2023). *E*.*coli* is preferred due to its simplicity, cost-effectivity, and widely-known genetic background (de Marco, 2025; Sezonov *et al*., 2014). On the other hand, some proteins can be produced in insoluble form because of limited responses of post-translational modifications in prokaryotic systems (Terpe, 2006; Nothaft & Szymanski, 2013; Walsh, 2010; Li *et al*., 2025; de Marco, 2025; Wang *et al*., 2023; Figueroa-Bossi *et al*., 2019; American Society for Microbiology, n.d.). For this reason, the outer and inner cellular environment optimizations have a pivotal role, especially for advanced structural biology techniques such as NMR that require isotopic labelling (Chae, Kim & Hyun, 2015; Chae & Kim, 2015; Chae, Kim & Um, 2019; Azatian *et al*., 2019; Wang *et al*., 2018).

M9 minimal media, initially originated from *Anderson et al*., serve a selective environment for isotopically labeled proteins (Anderson, 1946). However, the limited content of M9 generates metabolic stress for bacterial growth and may reduce the protein expression yield. The solutions to this problem have been developed by proposed M9+ and M9++ variants. *Cai et al*. presented the importance of low temperature conditions in minimal media by observing the increment in isotopic labeled protein expression yield compared to regular LB and M9 growing conditions (Cai *et al*., 2016). *Azatian et al*. highlighted the strong correlation between the increment in buffering capacity of M9 and higher protein expression level for highly deuterated 13CH3-side chain methyl group labeled samples (Azatian *et al*., 2019). In the same study, the developed M9++ reveals many advantages not only in cell densities but also in the yield of isotopic labeling usage (Azatian *et al*., 2019). For instance, the cell mass extracted from 250 mL M9++ cell culture is 50% greater than 1L M9 and requires 25% less 15NH4Cl usage (Azatian *et al*., 2019). In addition to these studies performed with *E*.*coli, Chae et al*. demonstrated the similar stressors’ effect, 6 different salt concentrations and 6 different pH levels, on *Saccharomyces cerevisiae* approximate 31 metabolites (Chae & Kim, 2015). Later, *Chae et al*. suggested that the recombinant protein expression in E.coli cells may rely on the relationship to cell metabolome profile investigation (Chae & Kim, 2015). The study hypothesized that different metabolite profiles can be linked to different protein production results in solubility, inclusion bodies or no production (Chae & Kim, 2015). To do this, they used 71 different genes and analyzed by several techniques including 2D 1-H-13C HSQC NMR and SDS-PAGE to identify the protein expression levels in soluble, inclusion bodies, and no production conditions (Chae & Kim, 2015). In total, only 21 metabolites are defined in the outer section of no production region suggesting in most cases metabolic profiling presents similarity (Chae & Kim, 2015). Among all metabolites, betaine, acetylphosphate, α-ketoisovalerate, and N-acetyllysine are found as inversely proportional to protein production. ^15^ These metabolites are determined as the highest level in no protein production and the least level in inclusion body cases (Chae & Kim, 2015). Besides, osmo-protection capability of betaine and its high level can diminish the expression and is linked to aggregation (Chae & Kim, 2015). Glutathione, on the other hand, is determined in the oxidized form which may be an indication of redox stress during the expression (Chae & Kim, 2015; Chae, Kim & Um, 2019). The studies for media enhancement and metabolic effects are identified separately and current literature information gives the media contents’ effect on protein expression level different from the media impact of metabolites. The optimized inner and outer shells of cells are not characterized in enhanced media conditions from metabolic identification perspective. Specially, M9+ and M9++ environments’ impact on metabolites during protein expression and the potential of cell stressors’ emergence are not fully understood. The lack of such information limits the carriage of cellular environment optimization to the advanced stage. Although Chae et al. identifies significant differences of metabolites between M9 variants are not evaluated specifically which is the novel emphasis of this study. In this study, outer cell metabolites are screened and compared for M9, M9+ and M9++ conditions during the protein expression by aiming to identify main stressor factors that limit the protein expression. Obtained metabolic entries will guide the development of new and optimized media as well as manipulate the metabolites which may potentially limit the protein expression. Overall, this study aims not only to increase the yield of expression level but also to enable background for more accessible and effective biological characterization via NMR spectroscopy *(Figure 1 and Figure 2)*.

**Figure 1.**
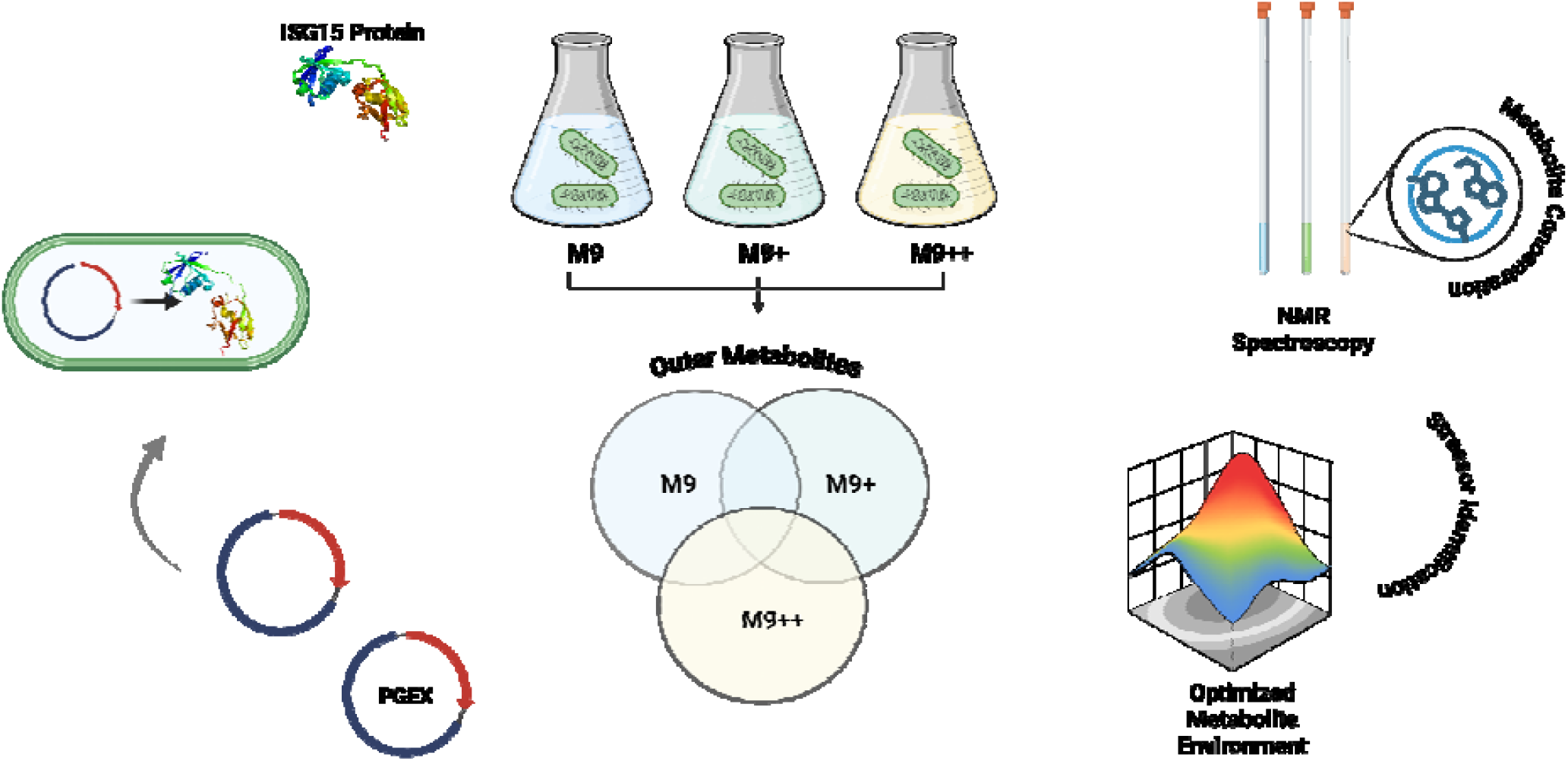
Workflow for optimizing metabolism of *E. coli* during ISG15 recombinant protein expression using M9 media formulations.

**Figure 2.**
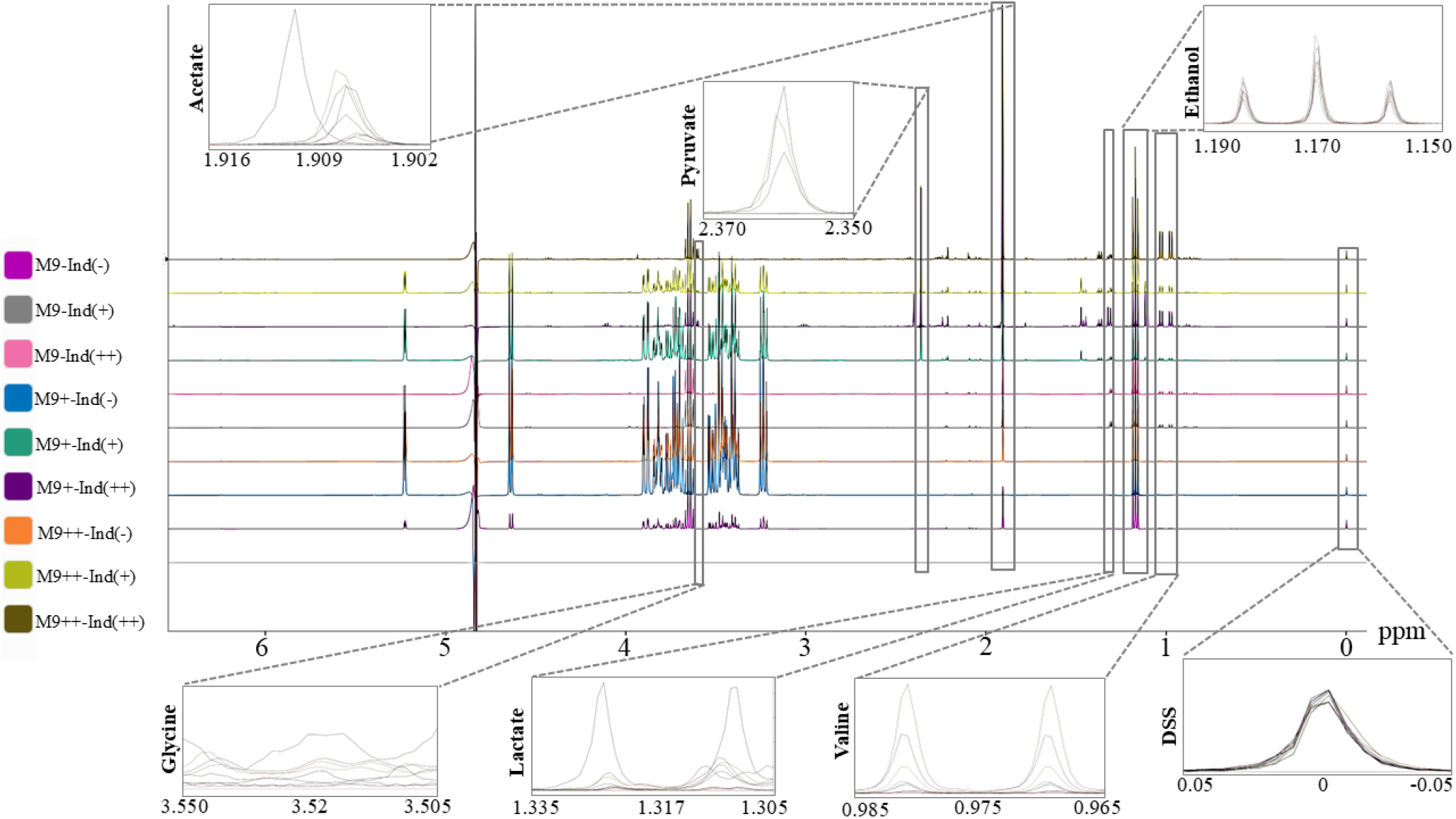
1D NMR analysis results and key metabolites demonstration for M9, M9(+), and M9(++) in induction (-), induction(+), and induction(++) conditions.

## Materials and Methods

### Plasmid Transformation and Starter Culture Preparation

Prior to initiating protein expression and large-scale bacterial culturing, a PGEX plasmid encoding the GST-fused ISG15 protein was transformed into *Escherichia coli* BL21 (DE3) competent cells. Transformation was performed using the standard heat-shock protocol with chemically competent cells. Following transformation, cells were plated onto LB agar supplemented with ampicillin (100 µg/mL) and chloramphenicol (25 µg/mL) and incubated overnight at 37□°C.

Single colonies were selected and inoculated into 25□mL of standard LB broth containing the same antibiotics. The LB medium was prepared by dissolving 10□g tryptone, 5□g yeast extract, and 10 g sodium chloride per liter of distilled water. The culture was incubated at 37□°C with shaking for at least 18 hours. After incubation, 750□µL of the bacterial culture was mixed with 750□µL of sterile glycerol in cryovials to prepare glycerol stocks for long-term storage at – 80□°C. These stocks were used to ensure experimental reproducibility from a single bacterial source.

### Preparation of LB and M9-Based Media

LB broth (BioBasic, Canada) was prepared by dissolving 25□g of commercially available LB powder in 1□L of distilled water. The solution was mixed until fully dissolved and then distributed into Erlenmeyer flasks in desired volumes (e.g., 25, 50, or 100□mL). Flasks were covered with aluminum foil and autoclaved at 120□°C for 90 minutes. The sterilized medium was stored at 4 °C until use.

For preparation of modified minimal media (M9+ and M9++) (Chae, Kim & Hyun, 2015; Chae, Kim & Um, 2019; Cai *et al*., 2016), the following components were dissolved in 97Lm□ of distilled water: K□HPO□, KH□PO□, Na□HPO□, K□SO□, D-glucose, and NH□Cl. The pH was adjusted to 7.2, and the solution was autoclaved at 120□°C for 90 minutes. After cooling, 1□mL of sterile-filtered 1□M MgCl□, 0.1□mL ampicillin, 0.1□mL chloramphenicol, and 1□mL of a vitamin solution (Sigma, USA) were added aseptically.

### Bacterial Growth and Media Transfer

Once the glycerol stocks and LB broth were prepared, an inoculum was prepared by transferring a glycerol stock into 50□mL of LB broth containing ampicillin and chloramphenicol. This culture was incubated at 37□°C with shaking for 15 hours. Following incubation, approximately 5□mL of culture was aliquoted into 1□mL portions in microcentrifuge tubes. The samples were centrifuged at 3000□rpm for 3 minutes, the supernatant was discarded, and the bacterial pellets were resuspended in 1□mL of sterile, pre-warmed M9+ or M9++ media. The resuspended cultures were then transferred into pre-warmed flasks containing 100□mL of the corresponding M9 medium. Optical density (OD_595_) was monitored regularly. When OD_595_ reached 0.3, a 2□mL sample was collected, and the incubation temperature was shifted to 22□°C to promote soluble expression of recombinant proteins. Samples were further collected at OD_595_ values of 0.6 and 0.8. When OD_595_ reached 0.8, IPTG was added to a final concentration of 0.4□mM to induce protein expression. Once OD_595_ reached 1.5, another 2□mL sample was collected, and the remaining culture was incubated at 22□°C for an additional 15 hours. At the end of the incubation, a final 2□mL sample was taken. All collected samples were centrifuged, and the supernatant and pellets were separated and stored at –80□°C for downstream analyses.

### NMR Sample Preparation and Data Acquisition

Frozen medium samples were thawed, and 500□µL of each was transferred into a tube. To each sample, 55□µL of a deuterium oxide-based buffer (37.7□mM Na□HPOL, 12.3□mM KH□POL, 20□mM NaCl, and 1□mM DSS) was added. DSS served as an internal chemical shift reference and was adjusted to a final concentration of 0.1□mM. Samples were transferred to 5□mm NMR tubes for data collection. NMR spectra were acquired on a 500□MHz Bruker Ascend spectrometer equipped with an Avance NEO console and a BBO room-temperature probe. The 1D NOESY-presat pulse sequence (noesygppr1d) was applied using 8K scans and 32K complex data points across a spectral width of 9615.4□Hz. Quantification of metabolite concentrations was performed using Chenomx NMR Suite 12 (Chenomx Inc., Canada), where DSS concentration and peak height were used as references. Eight thousand scans were applied to enhance the signal-to-noise ratio.

### Statistical Analysis of NMR Metabolomics Data

Metabolomics data were analyzed using MetaboAnalyst 6.0. Prior to statistical evaluation, the dataset was normalized using cube root transformation and Pareto scaling. Partial Least Squares Discriminant Analysis (PLS-DA) was employed to distinguish between time points and identify key metabolites contributing to temporal metabolic shifts. Variable Importance in Projection (VIP) scores from the PLS-DA model highlighted the most influential metabolites driving the observed metabolic changes throughout bacterial growth and recombinant expression.

## Results

### Effect of Media Composition on *E. coli* Growth Dynamics

To investigate the influence of nutrients on *E. coli* biomass accumulation during recombinant protein expression, bacterial cultures which contain the PGEX-GST-ISG15 plasmid were grown in three defined media conditions: standard M9, M9+, and M9++. Final optical density at 595 □nm (OD_595_) was recorded for each culture at the point of harvest, following IPTG induction and overnight incubation at 22□°C. All experiments were performed in biological replicates to ensure reproducibility. In the standard M9 minimal medium, the final OD_595_ values ranged from1.75 to 1.99, with a mean of 1.87□±□0.11(mean□±□SD), reflecting restricted growth under nutrient-limited conditions. The limited availability of carbon, nitrogen, and cofactors in M9 medium is consistent with the observed low cell densities, which can constrain recombinant protein yield.

In contrast, cultures grown in M9+ medium, which contains defined supplements including additional phosphate buffering, glucose, and trace vitamins, demonstrated markedly enhanced growth. The OD_595_ values for M9+ ranged from 5.88 to 6.45, with a mean of 6.21□±□0.24. This represents more than a threefold increase in cell density compared to M9, highlighting the critical role of nutrient availability in supporting bacterial biomass expansion during protein expression workflows. Interestingly, cultures grown in M9++ medium, which contains further supplementation beyond M9+, reached a slightly lower mean OD_595_ of 5.42□±□0.13. Although significantly higher than M9, the final cell densities in M9++ were moderately lower than those observed in M9+. This subtle reduction may indicate the onset of metabolic feedback inhibition or osmotic stress resulting from excessive nutrient concentration, which can modulate growth kinetics under semi-defined conditions.

Overall, the data suggest that moderate enhancement of M9 medium (M9+) promotes optimal *E. coli* growth for recombinant protein production, while further enrichment (M9++) does not confer additional benefit and may slightly hinder maximal growth. These findings underscore the importance of empirically optimizing medium formulations to balance metabolic support with physiological tolerance during heterologous protein expression in *E. coli*.

### Temporal Metabolite Profiling During Recombinant Protein Expression

To monitor dynamic metabolic changes during recombinant protein expression, samples were collected at three defined time points from each of the three biological replicates grown in M9, M9+, and M9++ media. These time points corresponded to 20 minutes prior to IPTG induction (pre-induction), the midpoint of the expression period(mid-expression), and at the end of the expression phase (15 hours post-induction). A total of 27 samples (3 media□×□3 replicates□×□3 time points) were analyzed by 1D ^1^H NMR spectroscopy. *(Figure 2)* After spectral acquisition, metabolite concentrations were quantified using Chenomx NMR Suite 12.00, employing DSS as an internal reference standard. This approach enabled absolute concentration determination for a panel of intracellular and extracellular metabolites. The NMR-based metabolic profiling revealed dynamic shifts in metabolite abundances across the expression timeline, which varied depending on the culture medium. Metabolomic analysis revealed distinct abundance profiles for key discriminatory metabolites among the three media conditions *(Figure 3 and Figure 5)*. Acetate was found at the highest concentrations in M9, intermediate in M9+, and lowest in M9++, suggesting that overflow metabolism is most pronounced under minimal nutrient conditions and progressively reduced with increased supplementation. Formate displayed an opposite trend, with the highest levels in M9++, lowest in M9+, and intermediate in M9, indicating enhanced mixed-acid fermentation activity in the most enriched medium. Acetoin was also most abundant in M9++, present at intermediate levels in M9+, and lowest in M9, highlighting the activation of alternative fermentation pathways under nutrient-rich conditions. In contrast, citrate reached its highest concentration in M9++, was intermediate in M9, and lowest in M9+, suggesting increased flux through the TCA cycle or altered citrate utilization as medium complexity increased (Zhang et al. 2019). Collectively, these patterns underscore the profound impact of media composition on central carbon metabolism during recombinant protein expression in E. coli.

**Figure 3.**
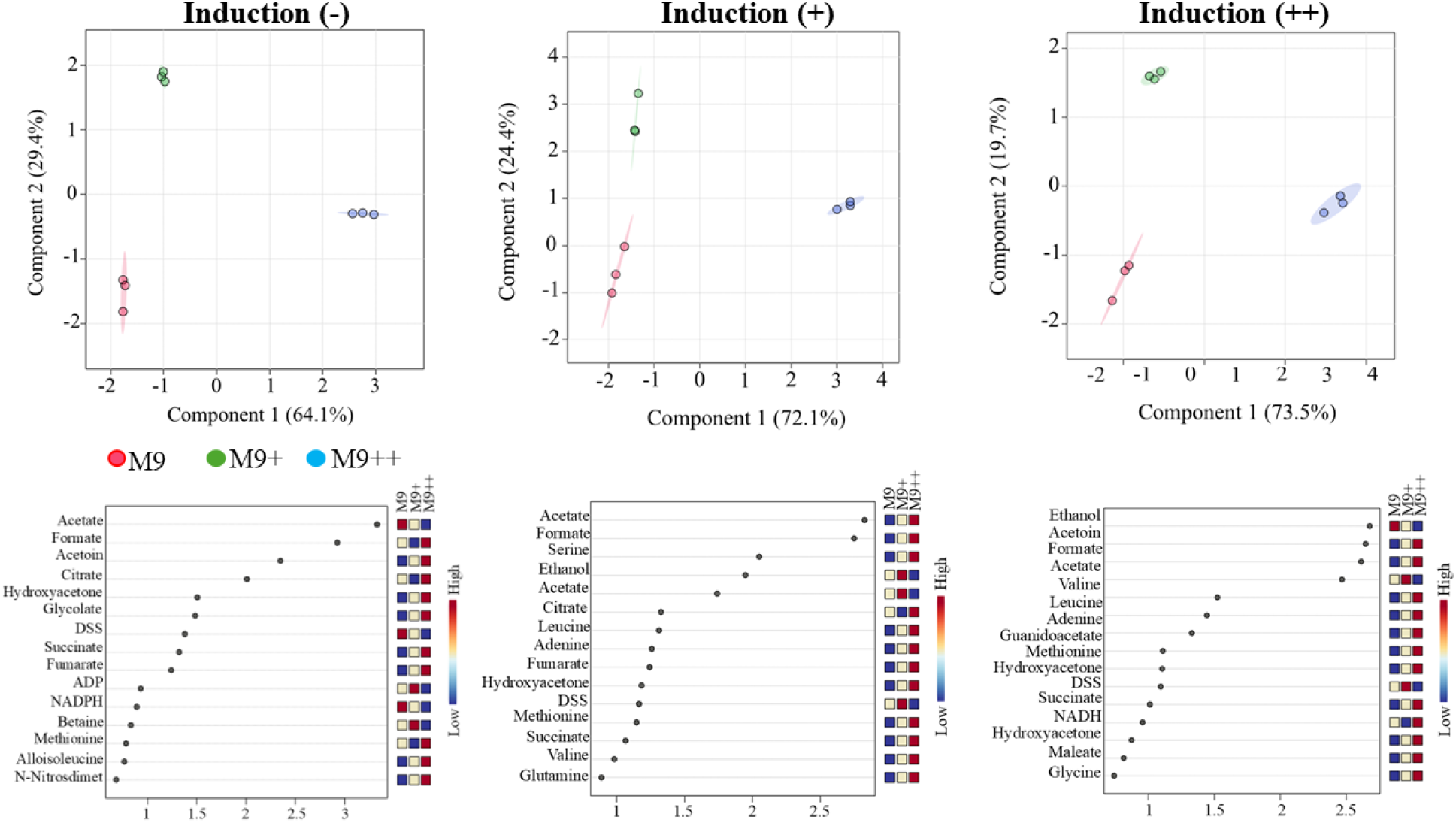
The comparison of PLS-DA (top) and VIP scores (bottom) of M9, M9+, and M9++ before induction (-), in induction (+), and further induction (++) without glucose among M9, M9+, and M9++ individually.

To further evaluate the significance of these metabolic changes, statistical analyses were performed using the MetaboAnalyst 6.0 platform. Comparative analyses focused on samples taken at the same time points across different media conditions, enabling the identification of metabolite variations specifically associated with media composition (*Figure 4*). PLS-DA was employed to distinguish the key metabolites between media groups, and VIP scores were used to identify key metabolites contributing to the observed differences. This analysis revealed that certain metabolites—such as acetate, ethanol, and amino acids such as serine—played prominent roles in differentiating metabolic responses among M9, M9+, and M9++ conditions. To specifically assess metabolic differences during active recombinant protein synthesis, a focused analysis was conducted on the mid-expression time point. At this stage, one sample from each of the three biological replicates was collected for each medium (M9, M9+, and M9++), yielding nine total samples for comparative evaluation. Multivariate statistical analysis using MetaboAnalyst revealed that metabolic profiles from M9+ and M9++ conditions clustered closely, indicating similar metabolite compositions under these enriched media. In contrast, M9 samples showed a pronounced separation from both M9+ and M9++, suggesting a distinct metabolic state driven by minimal nutrient availability. Among the statistically important metabolites, formate, serine, acetoin, and ethanol were identified as the top contributors to the observed separation, as determined by VIP scores from PLS-DA. Acetoin concentrations were the highest in M9+, intermediate in M9++, and lowest in M9, possibly reflecting the impact of initial glucose supplementation and its downstream utilization. Conversely, formate levels were lowest in M9, moderate in M9+, and highest in M9++. Serine followed a pattern similar to acetoin and formate, with the highest levels in M9++, intermediate in M9+, and lowest in M9, supporting the hypothesis of increased overflow metabolism in nutrient-rich environments.

**Figure 4.**
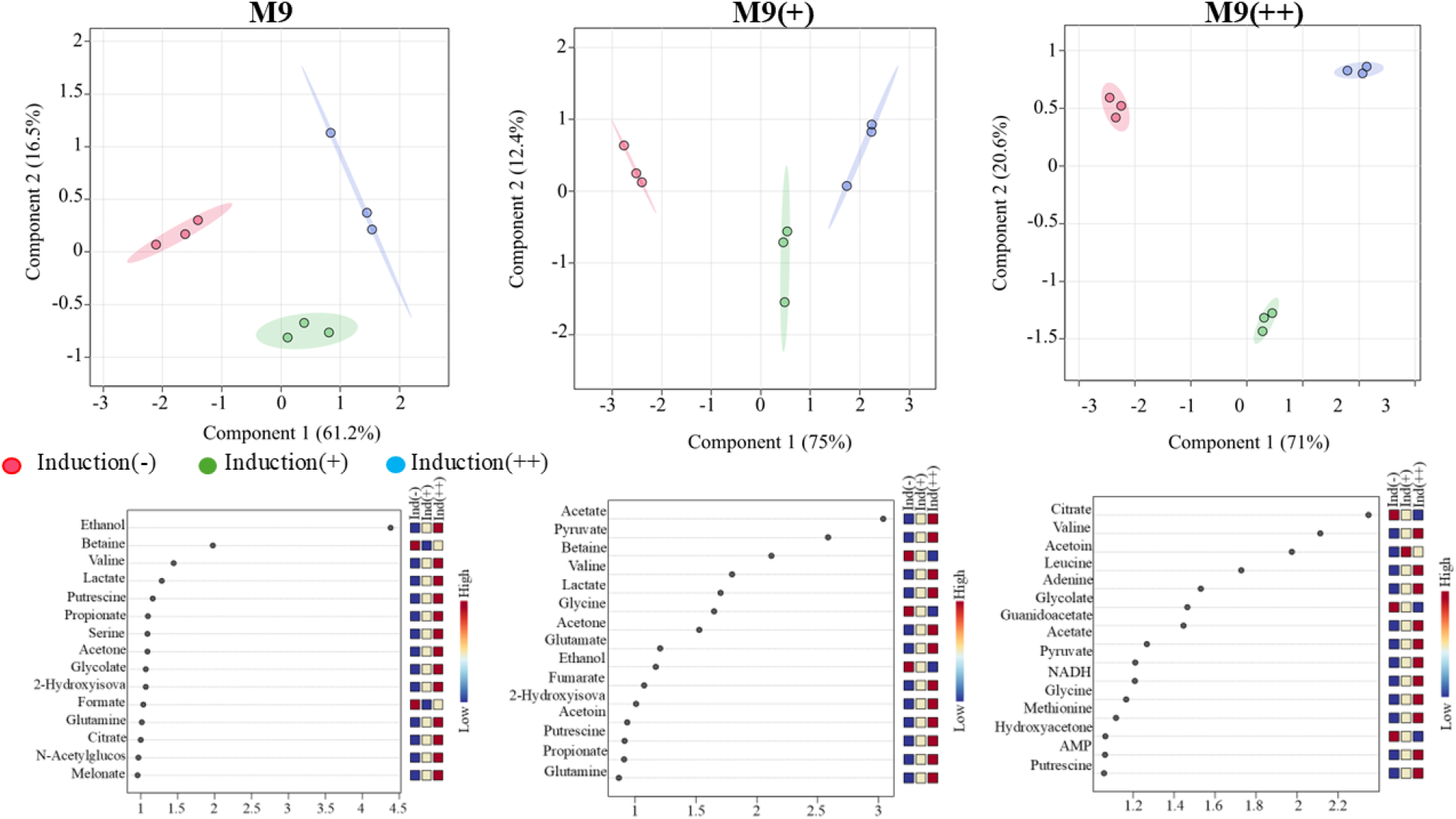
The comparison of PLS-DA (top) and VIP scores (bottom) of before induction (-), in induction (+), and further induction (++) among M9, M9+, and M9++ without glucose.

These results demonstrate that while M9+ and M9++ induce relatively similar metabolic states during expression, standard M9 medium results in a markedly different metabolic response, particularly in central carbon and fermentation-related metabolites. Such differences may have implications for protein yield, solubility, and overall cellular health during recombinant production *(Figure 5)*.

**Figure 5.**
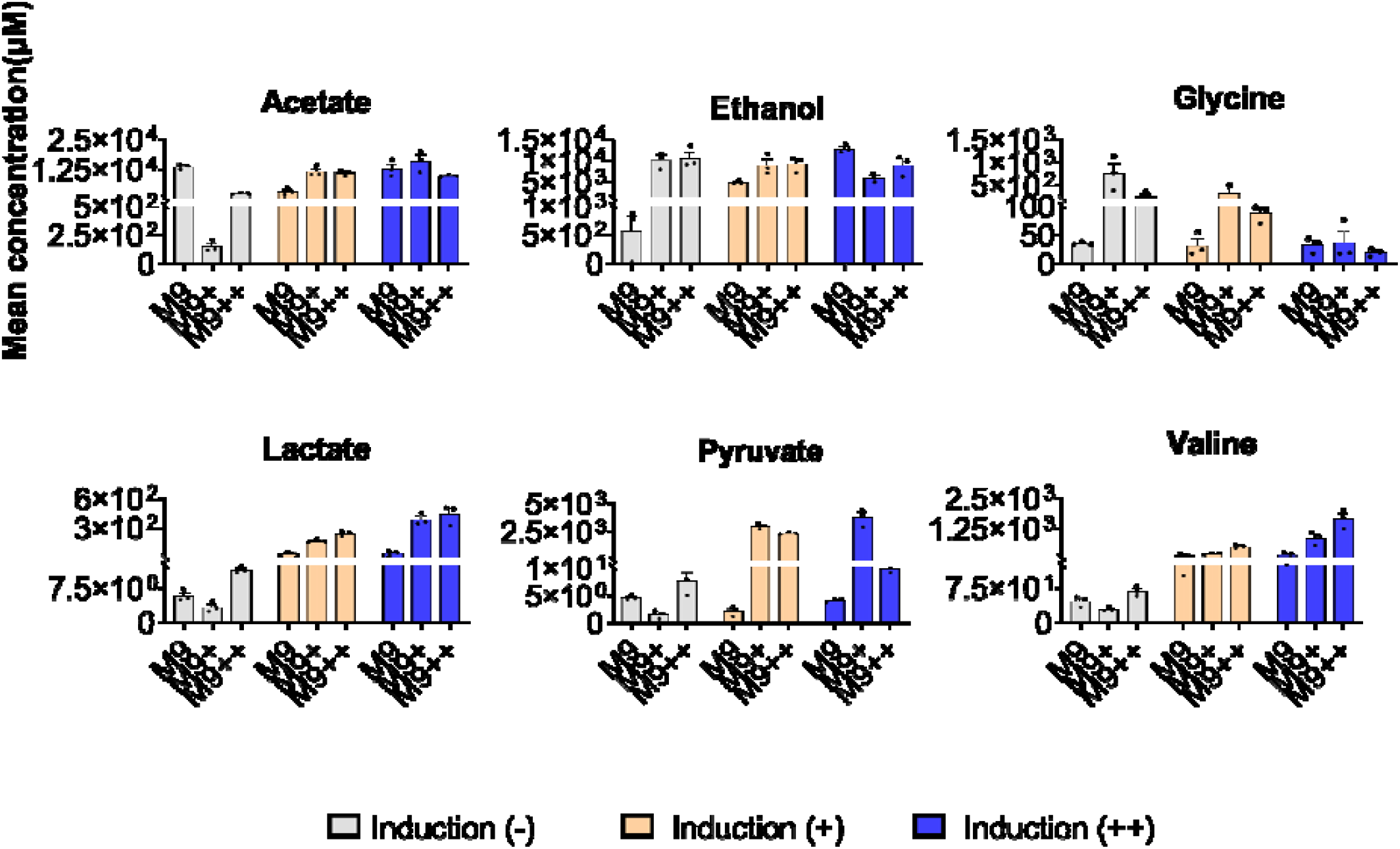
Key metabolites’ main concentration values in induction (-), induction (+) and induction (++) stages for M9, M9+ and M9++.

### Late-Stage Expression Reveals Divergent Amino Acid and Organic Acid Accumulation Across Media

At the final time point of recombinant expression (15 hours post-induction), NMR-based metabolomic analysis revealed significant media-dependent variations in central metabolic intermediates. Among the most discriminatory metabolites identified by VIP scores from PLS-DA were ethanol, acetoin, formate, acetate, and valine, each displaying distinct distribution patterns across the three media conditions. Ethanol levels were highest in cultures grown in M9, intermediate in M9+, and lowest in M9++, suggesting greater fermentative activity in nutrient-limited conditions and a shift toward more respiratory metabolism as nutrient supplementation increased. Conversely, both acetoin and formate exhibited the opposite trend, with highest concentrations observed in M9++, moderate levels in M9+, and the lowest in M9, indicating enhanced mixed-acid and overflow fermentation activity in the more enriched environments. Similarly, valine followed the same distribution as acetoin and formate, with the highest accumulation in M9++, moderate levels in M9+, and minimal levels in M9. This pattern suggests that valine biosynthesis is promoted under nutrient-rich conditions, likely due to increased precursor availability and energy supply. Acetate, however, demonstrated a distinct inverse trend. Its concentration was highest in M9+, moderate in M9, and lowest in M9++, indicating that overflow metabolism may be more pronounced in M9+ despite moderate nutrient availability, potentially due to imbalances in carbon flux and redox state.

These observations collectively indicate that media composition profoundly alters the metabolic state of *E. coli* during the late phase of recombinant protein expression. While M9 conditions drive fermentative pathways marked by elevated ethanol production, enriched media such as M9+ and M9++ shift metabolism toward enhanced biosynthesis and alternative fermentation end-products. Overall metabolites determined from induction and media conditions are demonstrated in *Table 1*. These metabolic reconfigurations likely reflect adaptive responses to differing nutrient landscapes and have implications for optimizing media formulations for improved protein yield and cellular health.

**Table 1.**
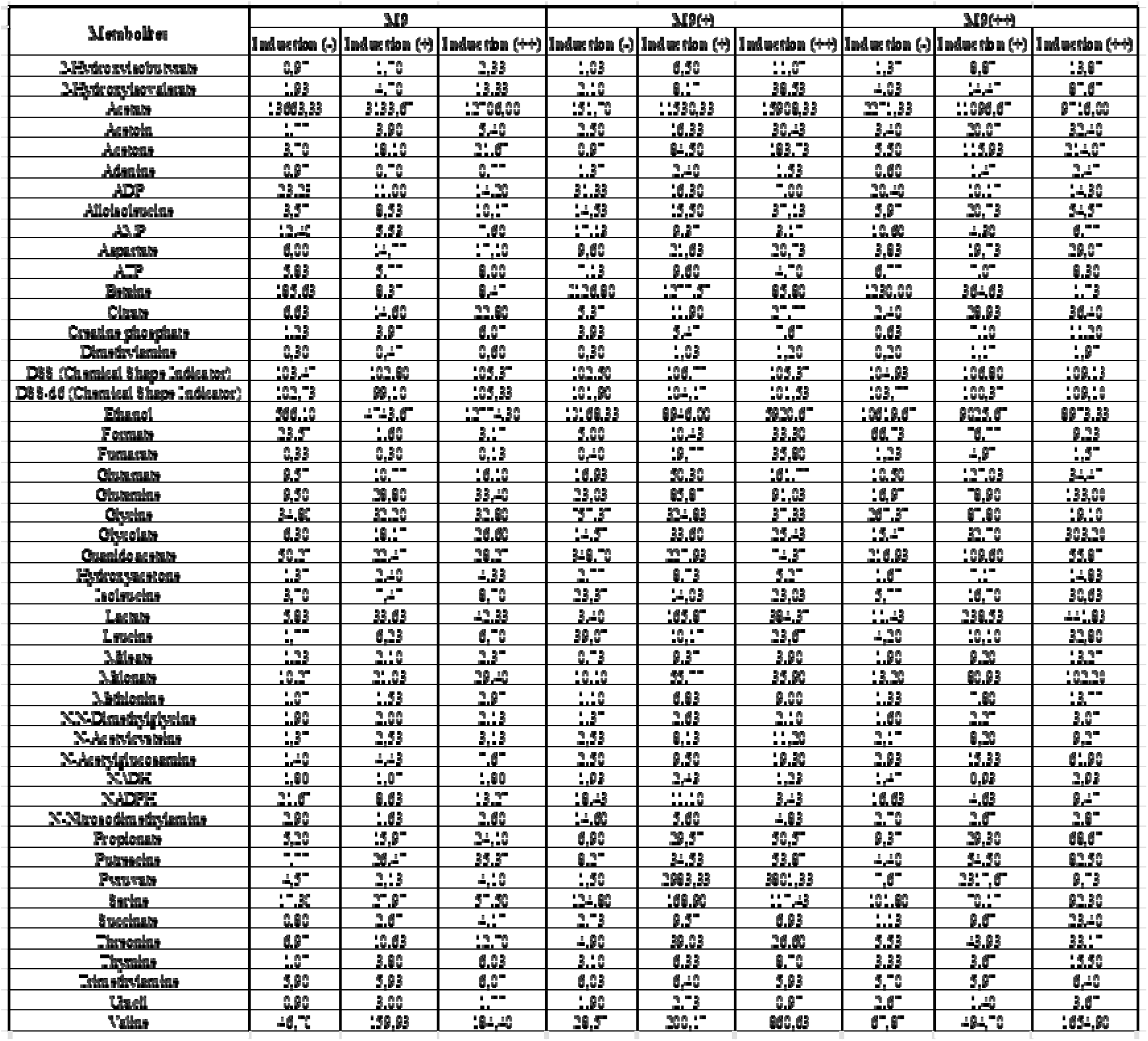
Overall metabolite concentrations (µM) for M9, M9+ and M9++ during recombinant protein expression (induction(-) (before induction), induction(+) (mid of expression, and induction(++)(end of expression)).

### Intra-Medium Metabolite Fluctuations During the Expression Timeline

In addition to comparing media types at specific time points, temporal metabolic changes within each medium—pre-induction, mid-expression, and post-expression—were analyzed to uncover the dynamic progression of metabolic states during recombinant protein production.

In M9 minimal medium, the metabolites most strongly contributing to separation across time points were ethanol, betaine, valine, lactate, and putrescine. Among these, Ethanol, valine, lactate, and putrescine exhibited a consistent increase over time, suggesting progressive activation of amino acid biosynthesis and stress-associated pathways. In contrast, betaine concentrations steadily declined over time, possibly reflecting its consumption as an osmoprotectant under prolonged culture conditions. In M9+ medium, the most significantly variable metabolites were acetate, pyruvate, valine, and lactate. Acetate, lactate, valine, and pyruvate concentrations increased over time, consistent with elevated metabolic flux and amino acid synthesis during active expression (Vemuri et al. 2006; Wolfe 2005). Conversely, glycine decreased over the expression period, suggesting a shift away from fermentative pathways and potential glycine utilization for protein synthesis or one-carbon metabolism. In M9++, the key metabolites distinguishing expression stages were citrate, valine, leucine, adenine. Valine, adenine and leucine showed a progressive increase over time, aligning with enhanced biosynthetic capacity in this enriched medium. In contrast, both citrate ethanol displayed a steady decrease, potentially reflecting metabolic adaptation to stable osmotic and redox environments (Chae & Kim, 2015). The results clearly demonstrate that media composition profoundly influences the metabolic state and growth dynamics of *E. coli* during recombinant protein expression. Specifically, standard M9 minimal medium, while advantageous for isotopic labeling due to its simplicity, significantly limits bacterial growth and induces a pronounced stress-related metabolic response. The elevated ethanol production observed in M9 indicates high fermentative activity, a common stress response due to nutrient scarcity and impaired respiration under minimal conditions. Although beneficial for specific labeling applications, such stress can negatively impact recombinant protein yield and solubility. In contrast, M9+ showed substantial improvement in bacterial growth and biomass accumulation. However, despite promoting high growth densities, M9+ displayed elevated acetate production, indicating pronounced overflow metabolism Enjalbert et al. 2015). Such overflow can reduce recombinant protein yield due to acidification of the medium and cellular toxicity, potentially limiting prolonged culture viability. Thus, while M9+ offers improved nutrient conditions compared to standard M9, its tendency toward overflow metabolism remains a critical limitation.

The M9++ medium, designed with enhanced buffering and nutritional supplementation, provided intermediate growth levels between M9 and M9+, with significant metabolic benefits. Reduced ethanol and acetate concentrations suggest a more balanced metabolic state and lower stress, allowing a shift toward biosynthesis of valuable metabolites such as valine, formate, and acetoin. These metabolites potentially reflect enhanced cellular fitness, which is crucial for maintaining higher-quality protein expression. Despite these advantages, the slightly lower growth compared to M9+ suggests potential osmotic or nutrient imbalance stress from excessive supplementation in M9++. Optimization of glucose concentration, buffering agents, and precise vitamin supplementation in M9++ may mitigate osmotic stress and further enhance bacterial performance. Additionally, adjustments to ammonium chloride and magnesium levels could optimize the balance between metabolic flux and cellular homeostasis, potentially improving overall recombinant protein yields.

## Conclusion

In conclusion, each medium presented distinct advantages and limitations. Standard M9 medium is optimal for isotopic labeling but severely restricts bacterial growth and induces metabolic stress. M9+ medium substantially enhances biomass but promotes overflow metabolism, potentially impacting protein quality and yield negatively. The M9++ medium represents an improved balance, reducing metabolic stress markers and promoting biosynthesis pathways beneficial for protein production. To further optimize M9++, targeted refinements should be considered to maximize its performance. First, glucose concentration should be carefully titrated to reduce overflow metabolism while ensuring an adequate carbon supply to support sustained growth. The buffering capacity of the medium can be fine-tuned by adjusting the ratios of phosphate salts to help stabilize intracellular pH without triggering phosphate repression. Finally, for large-scale or prolonged expression applications, controlled glucose feeding strategies can be employed to minimize acetate accumulation and support consistent protein yield.

Collectively, these optimizations have the potential to make M9++ a more robust and efficient medium tailored for high-quality recombinant protein expression in *E. coli*, particularly for applications requiring stable, isotopically labeled proteins for structural biology studies.

## ACKNOWLEDGMENT

CD acknowledges support from TUBITAK (Project No: 120Z594, 122Z747). OG acknowledges support from TUBITAK 2209 undergrad research support program. The authors acknowledge the use of the services and facilities of n^2^STAR-Koç University Nanofabrication and Nanocharacterization Center for Scientific and Technological Advanced Research. We gratefully acknowledge Kerem Kahraman for sample preparation.

## Author Contributions

**Ça**ğ**da**ş **Da**ğ: Conceptualization, Methodology, Supervision Funding acquisition, Investigation, Formal analysis, Writing-Reviewing and Editing, Writing-Original draft preparation **Oktay Göcenler:** Conceptualization, Methodology, Funding acquisition, Investigation, Formal analysis. **Nilüfer Çakır:** Statistical Analysis, Writing-Reviewing and Editing, Writing-Original draft preparation, Visualization. **Merve Turgut:** Investigation, Formal analysis. **Alp Eren Kazar:** Investigation, Formal analysis.

## References

American Society for Microbiology (ASM). LB (Luria-Bertani) agar protocol. https://asm.org/getattachment/5d82aa34-b514-4d85-8af3-aeabe6402874/lb-luria-agar-protocol-3031.html

Anderson, E. H. Growth requirements of virus-resistant mutants of Escherichia coli strain “B”. Proc. Natl. Acad. Sci. U.S.A. 32, 120–128 (1946).

Azatian, S. B., Kaur, N. & Latham, M. P. Increasing the buffering capacity of minimal media leads to higher protein yield. J. Biomol. NMR 73, 11–17 (2019).

Cai, M., Huang, Y., Yang, R., Craigie, R. & Clore, G. M. A simple and robust protocol for high-yield expression of perdeuterated proteins in Escherichia coli grown in shaker flasks. J. Biomol. NMR 66, 85– 91 (2016).

Chae, Y. K., & Kim, S. H. Searching for growth conditions for optimized expression of recombinant proteins in Escherichia coli by using two-dimensional NMR spectroscopy. Bull. Korean Chem. Soc. 36, 66–73 (2015).

Chae, Y. K., Kim, S. H. & Hyun, J. S. Probing metabolite space of Escherichia coli via growth medium composition as monitored by two-dimensional NMR spectroscopy. Chem. Biodivers. 12, 925–935 (2015).

Chae, Y. K., Kim, S. H. & Um, Y. Relationship between protein expression pattern and host metabolome perturbation as monitored by two-dimensional NMR spectroscopy. Bull. Korean Chem. Soc. 40, 634–641 (2019).

de Marco, A. Recent advances in recombinant production of soluble proteins in E. coli. Microb. Cell Fact. 24, 21 (2025).

Enjalbert B et al (2015) Acetate exposure determines the Diauxic behavior of Escherichia coli during the glucose-acetate transition. J Bacteriol 197(17):3173–3181

Figueroa-Bossi, N., Verdet, C. & Bossi, L. Bacteriophage–host interaction: Coordination between transcription and translation. Front. Microbiol. 10, 3096 (2019).

Li, J. et al. Detection and optimization of microbial expression systems for extracellular production and purification of Ca^2^D-responsive phase-changing annexin fusions. Protein Expr. Purif. 226, 106617 (2025).

Nothaft, H. & Szymanski, C. M. Bacterial protein N-glycosylation: New perspectives and applications. J. Biol. Chem.288, 6912–6920 (2013).

Overton, T. Recombinant protein production in bacterial hosts. Drug Discov. Today 19, 590–601 (2014).

Sezonov, G., Joseleau-Petit, D. & D’Ari, R. Escherichia coli physiology in Luria–Bertani broth. Front. Microbiol. 5, 172 (2014).

Terpe, K. Overview of bacterial expression systems for heterologous protein production: From molecular and biochemical fundamentals to commercial systems. Appl. Microbiol. Biotechnol. 72, 211–222 (2006).

Tiwari, S., Prakash, S. & Kumar, A. Comparative proteomics of Escherichia coli under different culture conditions reveals key metabolic pathway regulations. Sci. Rep. 14, 71238 (2024).

Vemuri GN et al (2006) Overflow metabolism in Escherichia coli during steady-state growth: transcriptional regulation and effect of the redox ratio. Appl Environ Microbiol 72(5):3653–3661

Wolfe AJ (2005) The acetate switch. Microbiol Mol Biol Rev 69(1):12–50

Walsh, G. Post-translational modifications of protein biopharmaceuticals. Drug Discov. Today 15, 773– 780 (2010).

Wang, H., Guo, J., Chen, X. & He, H. The metabolomics changes in Luria-Bertani broth medium under different sterilization methods and their effects on Bacillus growth. Metabolites 13, 958 (2023).

Wang, L., Liu, Q., Du, Y., Tang, D. & Wise, M. J. Optimized M9 minimal salts medium for enhanced growth rate and glycogen accumulation of Escherichia coli DH5α. Microbiol. Biotechnol. Lett. 46, 194– 200 (2018).

Zhang Y et al (2019) Pyruvate dehydrogenase E1 component regulates carbon metabolism and promotes growth and survival of E. coli under stress conditions. J Bacteriol 201(14):e00228–e00219

